## Supplemental figure for "LKB1 regulates JNK-dependent stress signaling and apoptotic dependency of *KRAS*-mutant lung cancers"

### Supplemental Figure Legends

**Supplemental Figure 1. Sensitivity of *KRAS$^{G12C}$*-mutant NSCLC cell lines to sotorasib and trametinib drug combinations**. **A.** Major co-occurring mutations in *KRAS*-mutant cell lines used in this study. **B.** Sotorasib sensitivity of *KRAS$^{G12C}$*-mutant NSCLC cell lines. Cells were treated with sotorasib for 3 days and viability was determined by CellTiter-Glo (CTG). **C.** Comparison of sotorasib sensitivity (quantified by AUC of sotorasib dose response curve) of *KRAS$^{G12C}$*-mutant cell lines grouped by co-occurring mutations. Each dot represents the mean AUC of 3 independent biological replicates. **D.** Calculation of ΔAUC as a metric of dependence on targeted alternate pathway in the presence of suppression of KRAS or MEK signaling. For instance, MCL-1 dependence as determined by relative sensitivity to sotorasib + AMG 176 compared to sotorasib alone. **E.** STK11 loss correlates with higher ΔAUC to sotorasib + AMG 176 and sotorasib + GDC-0941 in *KRAS$^{G12C}$*-mutant NSCLC cell lines. Cells were treated with sotorasib or sotorasib + AMG176/ navitoclax/ GDC-0941/ TNO155/abemaciclib for 3 days and viability was determined by CTG. **G.** ΔAUC of trametinib + AMG 176 (or the related compound AM-8621) versus trametinib alone. Each dot represents an independent biological replicate (N=3-4). **H.** LKB1-deficient cell lines are similarly sensitive to cobimetinib (MEKi) or adagrasib (KRAS G12Ci) combined with AMG 176. **I.** Single-agent AMG 176 sensitivity of *KRAS$^{G12C}$*-mutant NSCLC cell lines.

**Supplemental Figure 2. Inhibition of MCL-1 is synergistic with MAPK inhibition in cell lines with loss of LKB1 that exhibit high apoptotic responses. A.** Synergistic interaction between trametinib and AMG 176 in LKB1-loss cell lines. *KRAS$^{G12C}$*-mutant NSCLC cell lines were treated with increasing dose of trametinib and AMG 176 for 3 days and cell viability was determined by CTG. Synergy was calculated by the generalized Loewe method (see Method). Data are averaged from 3 independent biological replicates for each cell line. **B.** Cytotoxicity was observed in *LKB1*-deficient *KRAS$^{G12C}$*-mutant NSCLC cell lines with combined MAPK + MCL-1 inhibition. *KRAS$^{G12C}$*-mutant NSCLC cell lines were treated with increasing doses of trametinib/sotorasib + AMG 176 for 3 days and viability was determined by CTG. Growth Rate Inhibition was calculated by normalizing to vehicle treated cells (0), day 0 cell viability (100), and no viable cells (200) using the same data as synergy calculation in Panel A.

**Supplemental Figure 3. LKB1-deficient cells have increased sensitivity to combined MAPK + MCL-1 inhibition. A.** Left: Expression of LKB1 in LKB1-deficient *KRAS*-mutant NSCLC cell lines. (EV = pBABE empty vector control). Right: CRISPR-mediated knockout of LKB1 in LKB1-deficient *KRAS*-mutant NSCLC cell lines. (KO GFP = sgGFP negative control). **B.** Relative proliferation of isogenic cell line pairs. **C-D.** Dose response curves for H2030 EV, H2030 LKB1, H358 KO GFP, and H358 KO LKB1 cell lines after treatment with increasing doses of trametinib/sotorasib in the absence or presence of 1 µM of AMG 176. Data is representative of three independent biological replicates (N=3). **E.** Re-expression of LKB1 in LKB1-deficient cells, or LKB1 deletion in LKB1 WT cells, does not alter sensitivity to sotorasib alone (expressed as AUC of sotorasib dose response curve). Each dot represents an independent biological replicate. **F.** Restoration of LKB1 in LKB1-deficient *KRAS*-mutant cell lines, or LKB1 deletion in LKB1 WT cells, increases apoptotic response (annexin + cells) to trametinib + AMG 176, assessed by live cell imaging. Data are mean and S.E.M. of 3 technical replicates. **G.** Cytotoxic versus cytostatic response as determined by calculating Growth Rate Inhibition (vehicle treated cells = 0, day 0 cell viability = 100, no viable cells = 200). Cells were treated for 3 days and viability determined by CTG. Data are mean and S.E.M. from 3 independent biological replicates. **H.** Waterfall plots showing degree of tumor volume change of H2030 isogenic xenograft tumors after treatment with sotorasib (30 mg/kg daily), trametinib (3 mg/kg daily), or in combination with AMG 176 (50 mg/kg daily) for 3.5 weeks.

**Supplemental Figure 4. Phosphoproteomic analysis reveals JNK activation in LKB1-deficient *KRAS*-mutant cells after drug treatment. A.** Quantification of $\Delta$AUC from CTG assays of *KRAS*-mutant NSCLC cell lines with LKB1 loss (EV), LKB1 restoration (LKB1), or restoration of a kinase dead version of LKB1 (LKB1$^{K87I}$ kd) after treatment with trametinib + 1 µM AMG 176. Each dot represents an independent biological replicate (N=3). **B.** Western blot assessment of LKB1 and pAMPK (T172) in H2030 EV, H2030 LKB1, and H2030 LKB1 kd cells after treatment with 1 mM AICAR for 8 hours. **C-D.** Differential levels of phosphopeptides between trametinib and vehicle treatment in H2030 EV, H2030 LKB1, and H2030 LKB1 kd cell lines (grouped). Number of differentially quantified phosphopeptide and proteins with fold-change > 2-fold and adjusted p value < 0.05 are indicated. **E.** Volcano plots showing differentially enriched signatures between trametinib and vehicle treatment. Phosphopeptide signatures were calculated using ssGSEA2.0/PTM-SEA. **F.** Differentially enriched phosphopeptide signatures in paired isogenic cell lines in the absence of drug treatment. **G.** Differential enrichment of phosphopeptide signatures in trametinib + AMG 176-treated isogenic cell line pairs. **H.** Differential phosphorylation of individual JNK substrates in trametinib + AMG 176 treated isogenic cell line pairs.

**Supplemental Figure 5. JNK activation in LKB1-deficient cells underlies MCL-1 dependence. A.** Western blot analysis of isogenic *KRAS*-mutant NSCLC cell lines after treatment with 1 µM sotorasib + 1 µM AMG 176 (SA) or 0.1 µM trametinib + 1 µM AMG 176 (TA) for 8 hours. **B.** H2030 cells were treated with sotorasib/trametinib + AMG 176 or UV irradiation for up to 4 hours and phospho-JNK was assessed by immunofluorescence. **C.** Left: Effect of siRNA knock-down of MKK4 and MKK7 on pJNK in H2030 EV cells treated with 0.1 µM trametinib (T) for 24 h or 0.1 µM of trametinib for 24 h followed by 1 µM AMG 176 (TA) for 4 hours. Right: mRNA expression of MKK4 and MKK7 after siRNA knockdown was determined by RT-qPCR. **D.** H2030 isogenic cell lines with empty vector or LKB1 expression were treated with sotorasib + AMG 176 for up to 8 hours and phospho-JNK (left) and annexin staining (right) were performed to assess the dynamics of JNK activation and apoptosis, respectively. **E.** JNK phosphorylation in *KRAS*-mutant NSCLC cell lines treated with 0.1 µM of trametinib + 1 µM of AMG 176 (TA) for 8 hours. **F.** *KRAS*-mutant NSCLC cell lines were treated 0.1 µM trametinib + 1 µM AMG 176 for 8 hours. Data on X axis show quantification of densitometry levels from western blots of phospho-JNK normalized to total JNK, in drug-treated compared to vehicle cells from western blots. Data on Y axis show relative sensitivity to trametinib + AMG 176. Data are from 3 independent biological replicates for each cell line, Spearman correlation r=0.75, p=0.0331). **G.** Time-course of JNK phosphorylation in H23 EV and LKB1 cells treated with 0.1 µM trametinib + 1 µM AMG 176. **H.** JNK phosphorylation in H2030 EV and LKB1 cells after UV irradiation for 0-4 hours. **I.** JNK phosphorylation in H2030 EV cells with siRNA knockdown of JNK1+2 after treatment with 0.1 µM of trametinib for 24 hours (T) or 0.1 µM of trametinib for 24h + AMG 176 for 4 hours (TA). **J.** H2030 EV cells with siRNA knockdown of JNK1+2 (or siNC negative control) or H2030 LKB1 cells were treated with trametinib or sotorasib alone or in combination with 1 µM of AMG 176. Cell viability was determined by CTG after 3 days. Data is representative of 3 biological replicates. **K**. H2122, H23, LU65 EV cells with siRNA knockdown of JNK1+2 (or siNC negative control) were treated with trametinib + AMG176 for 48 hours and apoptosis induction (annexin positivity) was measured by live-cell imaging. Data are mean and S.E.M. of 3 technical replicates.

**Supplemental Figure 6. Suppression of JNK activation by LKB1 is mediated by NUAK kinases. A-D.** siRNA knockdown of AMPK-family assessed by western blot or RT-qPCR. **E-F.** Viability assessment H2030 LKB1 cells with corresponding siRNA knockdown after treatment with sotorasib/trametinib alone or in combination with 1 µM AMG 176. Data are representative of 3 biological replicates. **G.** Viability assessment of H2030 EV or LKB1 cells in culture media containing no glucose, 0.2 g/L glucose, or 2 g/L glucose. Data are representative of 3 biological replicates. **H.** LKB1-reconstituted (H2030, MGH1112) or LKB1 WT (H358) with siRNA knockdown of NUAK1+2 were treated with trametinib + AMG 176 and apoptosis (annexin positivity) was measured by live cell imaging. Data are mean and S.E.M. of 3 technical replicates. **I.** Viability assessment H2030 EV and LKB1 cells with siRNA knockdown of PP1B after treatment with sotorasib/trametinib alone or in combination with 1 µM AMG 176. Data is representative of 3 biological replicates.

**Supplemental Figure 7. Related to Figure 3. LKB1 deficiency increases BIM:MCL-1 interaction and creates an MCL-1 dependent state. A.** Schematic of BH3 profiling experimental setup. The change in priming (ΔPriming) is measured before and after treatment with trametinib (0.1 µM) as depicted in Figure S7C. **B.** Overall priming of *KRAS*-mutant NSCLC cells treated with vehicle or 0.1 µM of trametinib for 16 hours, as determined by BH3 profiling with titration of BIM BH3 peptide. **C.** Illustration of method for calculating Δpriming (increase or decrease in priming between vehicle and drug treated cells) from BH3 profiling results. **D-E.** Change in MCL-1 or BCL-XL dependence (MS1 BH3 peptide at 30 µM dose, HRK BH3 peptide at 100 µM dose) upon treatment of isogenic cell lines with trametinib. **F.** Expression of MCL-1 and BCL-XL in *KRAS*-mutant NSCLC cell lines grouped according to sensitivity to trametinib/sotorasib + MCL-1 inhibitor. **G.** Relative protein expression level of MCL-1 and BCL-XL in *KRAS*-mutant NSCLC cell lines grouped by LKB1 status from CCLE proteomic database.

**Supplemental Figure 8. LKB1 deficiency correlates with BIM:MCL-1 protein binding. A.** Method for calculating BIM:MCL-1 and BIM:BCL-XL binding ratios from co-immunoprecipitation (co-IP) of *KRAS*-mutant NSCLC cell lines. Input and IP protein bands were quantified from the same blot membrane. **B.** Co-IP assessment of BIM bound to MCL-1 and BCL-XL binding in *KRAS*-mutant NSCLC cell lines after treatment with 0.1 µM of trametinib for 24 hours. **C.** Quantification of BIM bound to MCL-1 versus BCL-XL in *KRAS*-mutant NSCLC cells after treatment with trametinib. BIM:MCL-1 and BIM:BCL-XL binding ratios were calculated from densitometry measurements as described in Fig. S8A. Input and IP protein bands were quantified from the same blot. **D.** Method for calculating BIM: MCL-1 and BIM-BCL-XL binding ratios in isogenic cell lines. Input and IP protein bands were quantified from the same blot membrane. Blot is reprinted from Fig. 3A for illustration purposes. **E-F.** Co-IP assessment of BIM bound to MCL-1 in isogenic cell lines after treatment with 0.1 µM of trametinib for 24 hours (T), or 0.1 µM of trametinib for 24 hours + 1 µM of AMG 176 for 4 hours (TA). **G.** BIM:MCL-1 binding ratios after 24 hours trametinib treatment (left) or vehicle (right) in isogenic cell lines. Binding ratios were calculated from densitometry measurements as shown in Fig. S8D.

**Supplemental Figure 9. JNK phosphorylates BCL-XL S62, altering BIM:BCL-XL binding and driving an MCL-1 dependent state. A-B.** Time-course of phosphorylation of MCL-1 and BCL-XL in H2030 EV/LKB1 and H358 KO GFP/LKB1 cells treated with 0.1 µM of trametinib. Blots shown in each panel are from a single membrane with non-relevant intervening lanes removed (indicated by solid vertical line). **C.** JNK1/2 knockdown in H2030 EV cells decreases drug-induced BCL-XL phosphorylation to a similar level as in H2030 LKB1 cells. After siRNA transfection, cells were treated with 0.1 µM trametinib for 48h or trametinib for 48h followed by 1 µM AMG 176 for 4 hours. **D.** Schematic of phospho-sites in MCL-1 and BCL-XL phosphorylated by JNK. MCL-1 E125 is a phospho-mimetic site, indicated in red. **E.** Western blot of H2030 EV cells with inducible WT MCL-1-Flag and BCL-XL-HA cultured in media containing various concentrations of doxycycline (DOX). **F.** Western blot of H2030 EV cells with inducible BCL-XL WT and siRNA knockdown of BCL-XL (or negative control) cultured in media containing various concentrations of DOX. **G.** MCL-1 phospho-mutants lack corresponding phosphorylation bands. **H.** Viability assessment of H2030 LKB1 cells with corresponding siRNA knockdown after treatment with sotorasib/trametinib alone or in combination with 1 µM AMG 176. Data are representative of 3 biological replicates. **I.** Viability of H2030 cells reconstituted with DOX-inducible BCL-XL WT or S62A mutants was assessed without DOX in the cell culture medium. Data are representative of 3 biological replicates. **J.** Viability of H2030 cells reconstituted with DOX- inducible BCL-XL WT or S62A mutant was assessed with optimal DOX concentration to restore endogenous levels as identified in Fig. S9E. Data are representative of 3 biological replicates. **K.** The BCL-XL S62A mutant reduces sensitivity of LKB1-deficient cell lines to trametinib or sotorasib + AMG 176. BCL-XL constructs were induced in H2122, H23 or MGH1112-1 cells and sensitivity to trametinib or sotorasib + AMG 176 was measured by CTG. Each dot represents an independent biological replicate (N=3). **L.** Co-IP assessment of BIM bound to BCL-XL WT or S62A in MGH1112 EV cells after treatment with 0.1 µM trametinib for 24 hours followed 1 µM AMG 176 for 4 hours.

**Supplemental Figure 10. BH3 profiling of *ex vivo* treated patient *KRAS*-mutant NSCLC cells and *in vivo* treated PDX tumors. A.** *KRAS*$^{G12C}$-mutant NSCLC tumor cells from patient metastatic sites were treated ex vivo with sotorasib or trametinib for 16 hours. Overall priming is shown, as determined by BH3 profiling with titration of BIM BH3 peptide. Change in BIM dependent priming of tumor cells after ex vivo treatment with 0.1 µM trametinib or 1 µM sotorasib treatment compared. Each dot represents a technical replicate. **B.** Mice bearing *KRAS*$^{G12C}$-mutant NSCLC PDX tumors were treated with trametinib (3 mg/kg) for 3 days and harvested for BH3 profiling. Data shown is overall priming as determined by titration of BIM BH3 peptide. Each dot represents an independent tumor, 5-10 animals were used for each treatment group/model.

**Supplemental Figure 11. *In vivo* response of *KRAS*-mutant NSCLC PDX models to sotorasib + AMG 176. A.** Waterfall plots showing tumor response of *KRAS*$^{G12C}$-mutant NSCLC PDX tumors treated with sotorasib (100 mg/kg daily), AMG 176 (50 mg/kg daily), or combination of sotorasib + AMG 176 at indicated time points. **B.** Animal body weights of PDX models treated with sotorasib, AMG176 or combination. 7-13 animals were used for each treatment group/model. **C.** Survival curves of PDX tumors: the progression free survival (PFS) metric was determined by time to 20% increase in tumor volume from baseline measurement. **D.** Mice bearing MGH1196-2 PDX tumors were treated with sotorasib (100 mg/kg) daily and AMG 176 (50 mg/kg) twice weekly. Tumor response was similar to AMG 176 dosed daily (see Figure 5E). Data are mean and S.E.M., N=4 animals per treatment group/model. **E-F.** Humanized MCL-1 knock-in mice were treated with sotorasib (100 mg/kg) daily with twice weekly dosing of AMG 176 (50 mg/kg). Body weight was monitored. B cells, T cells, plasma cells, monocytes, and neutrophils from peripheral blood (24 hours after cycle 2) were characterized by flow cytometry. Each dot corresponds to an individual mouse (N=4 per treatment group).

Supplemental Table 1. IC50 and EMAX of sotorasib in KRAS NSCLC cell lines.

Supplemental Table 2. RT-PCR primer sequences used in this study

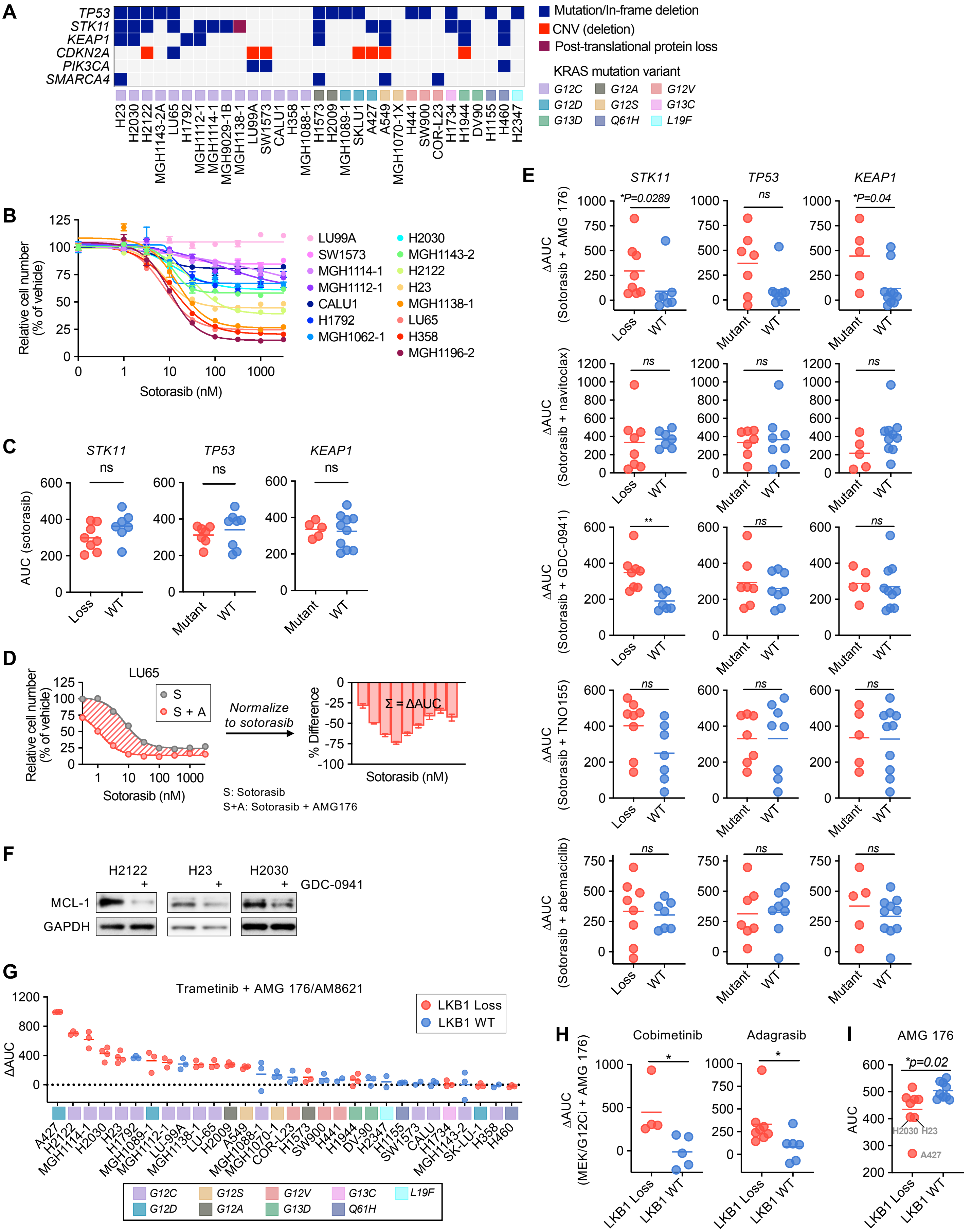


Supplementary Figure 1

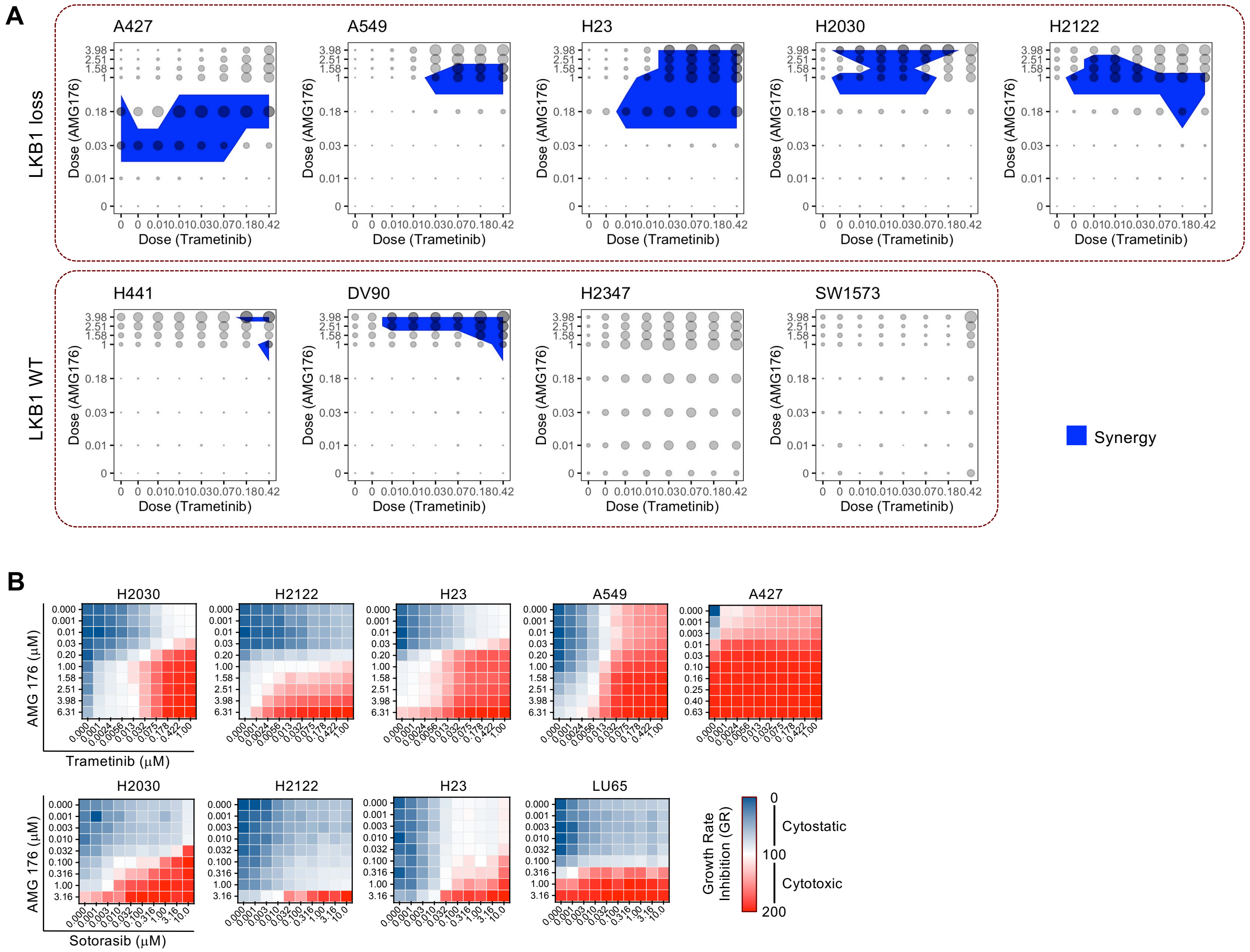


Supplemental Figure 2

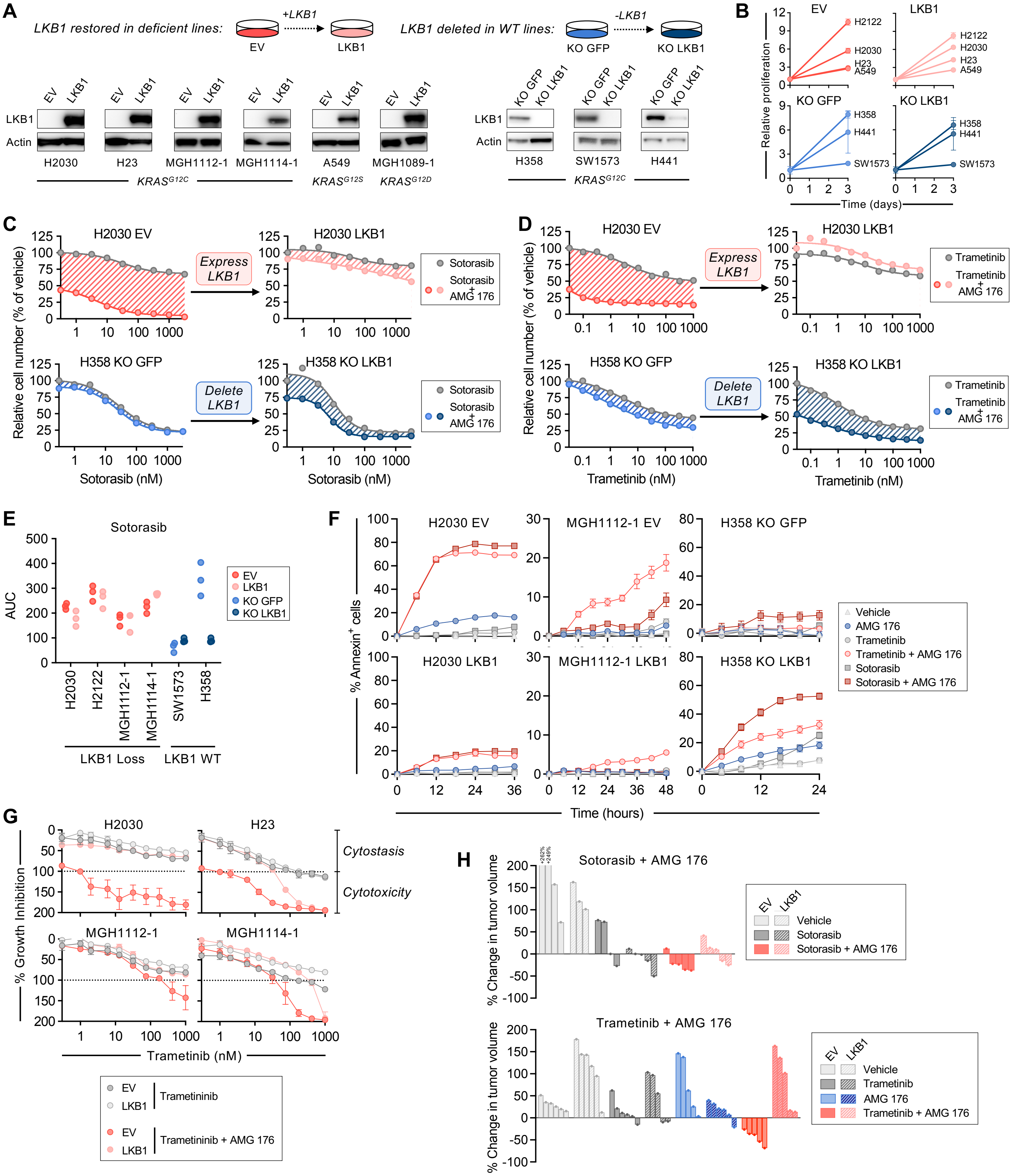


Supplemental Figure 3

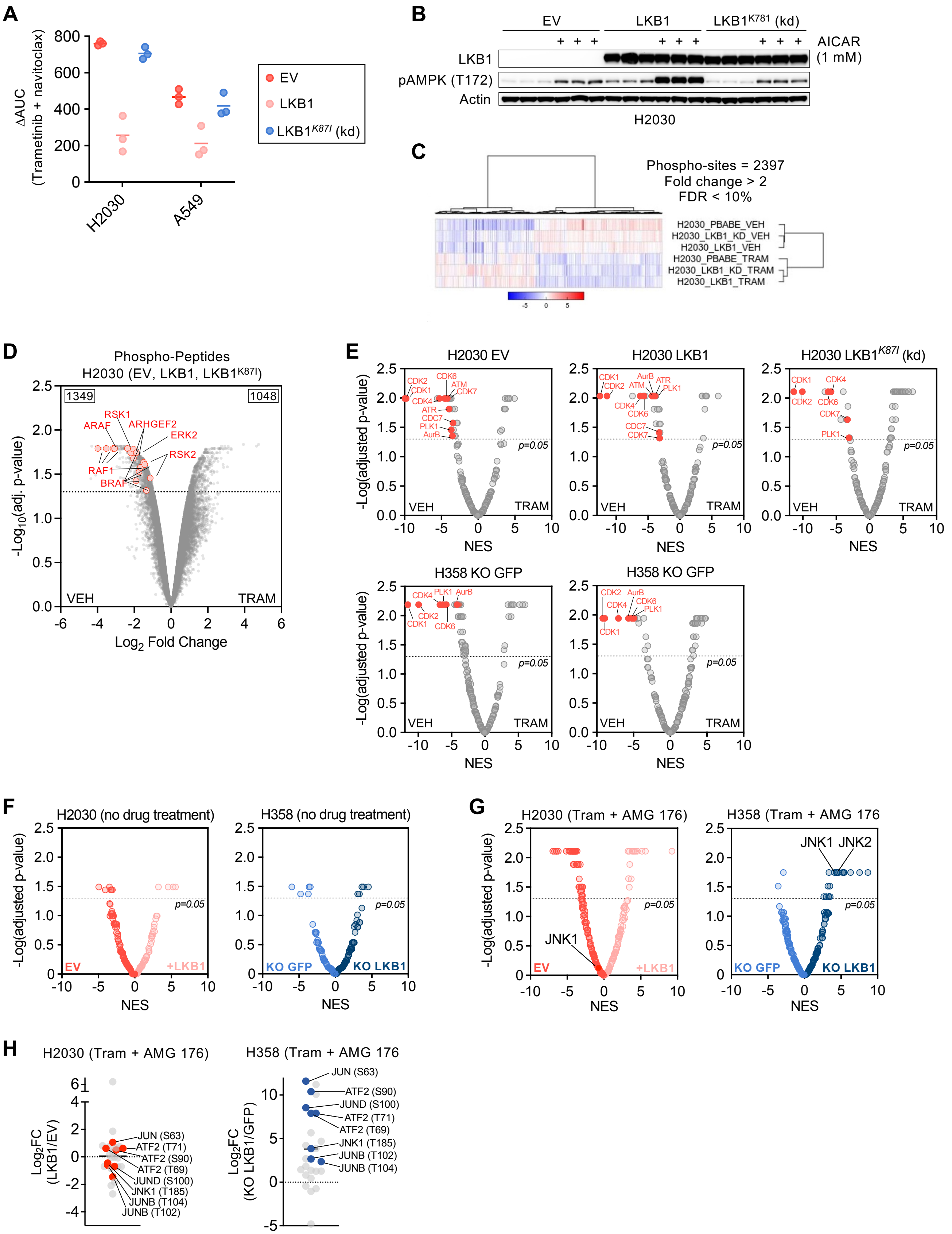


Supplemental Figure 4

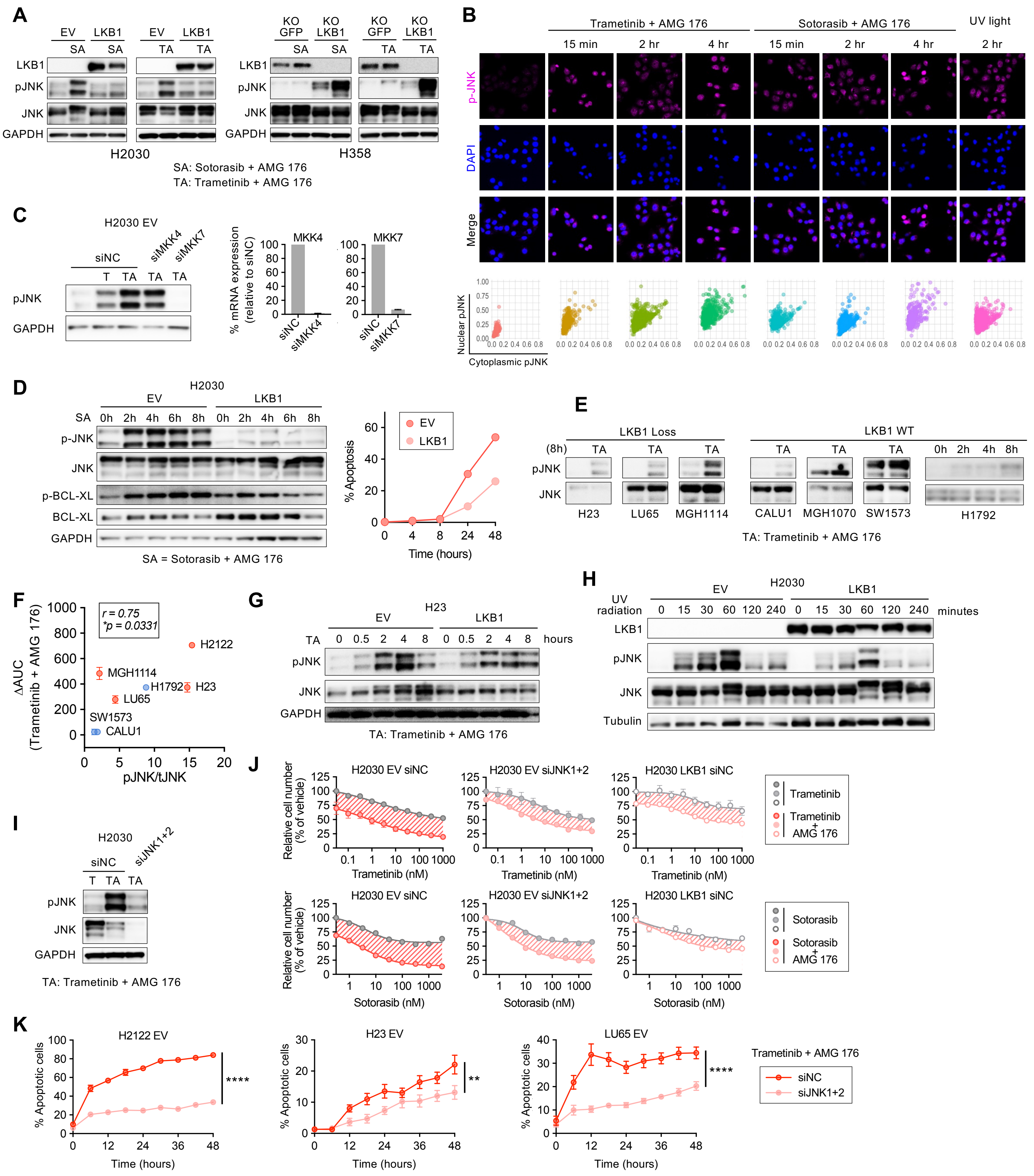


Supplemental Figure 5

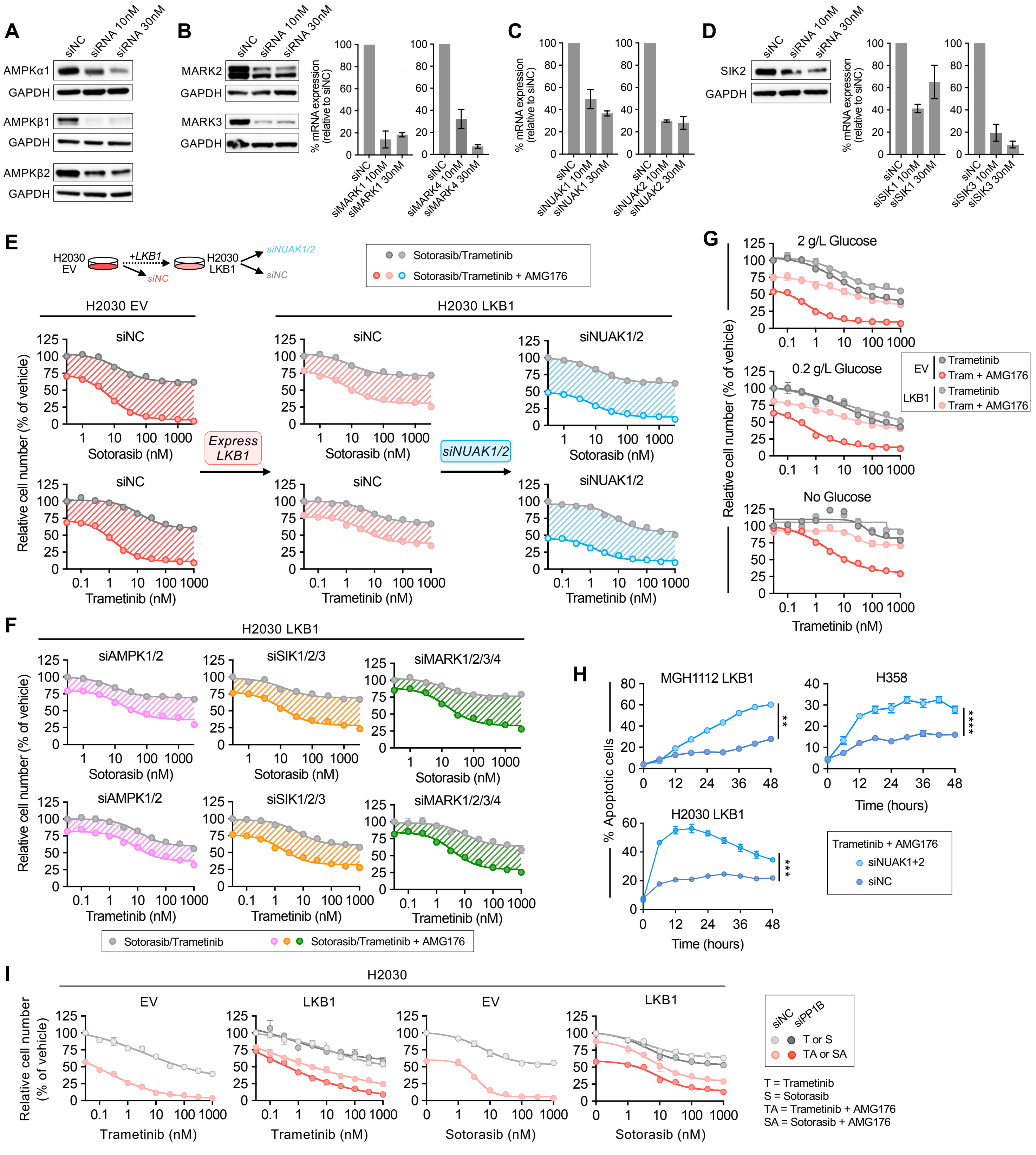


Supplemental Figure 6

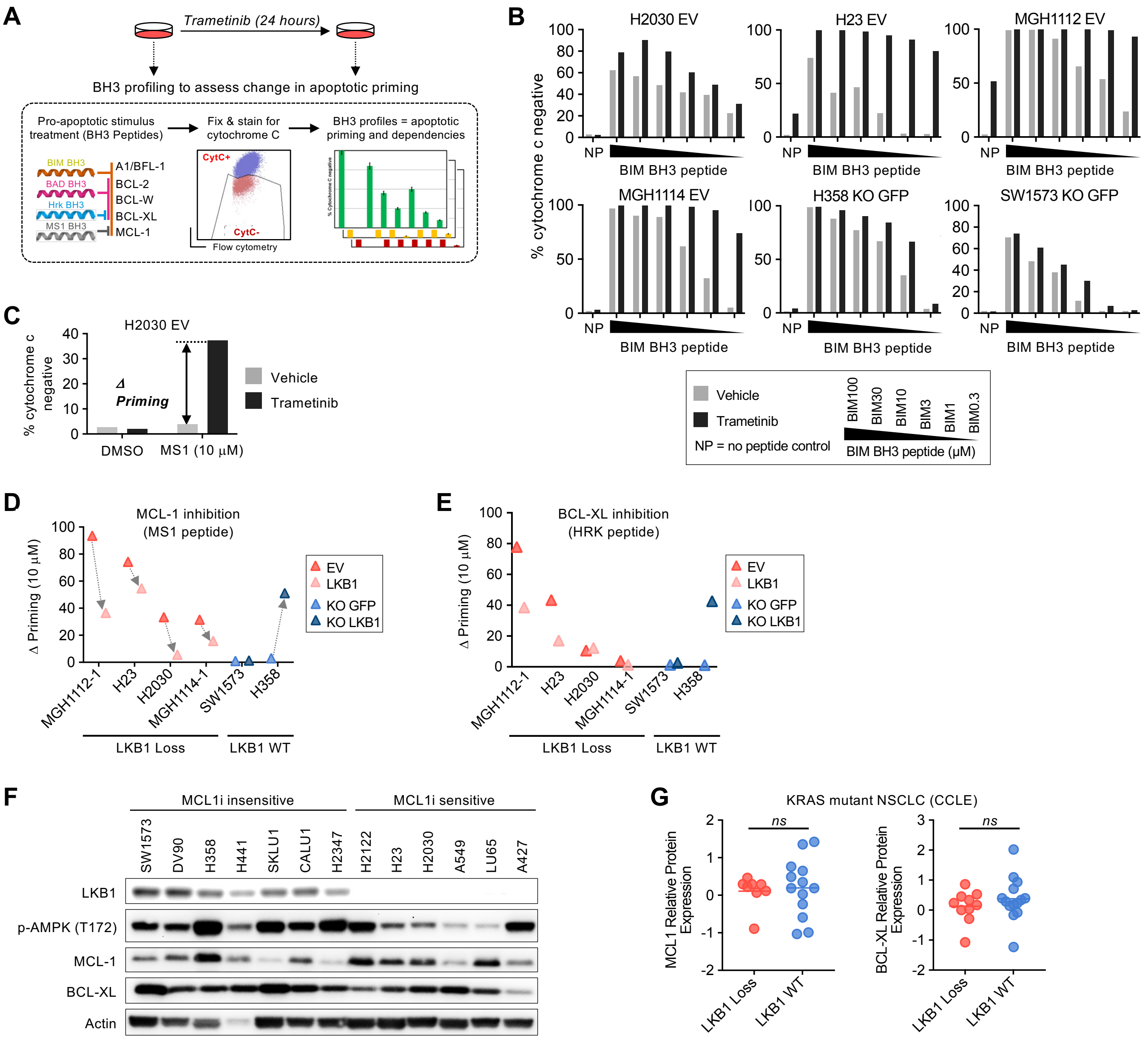


Supplemental Figure 7

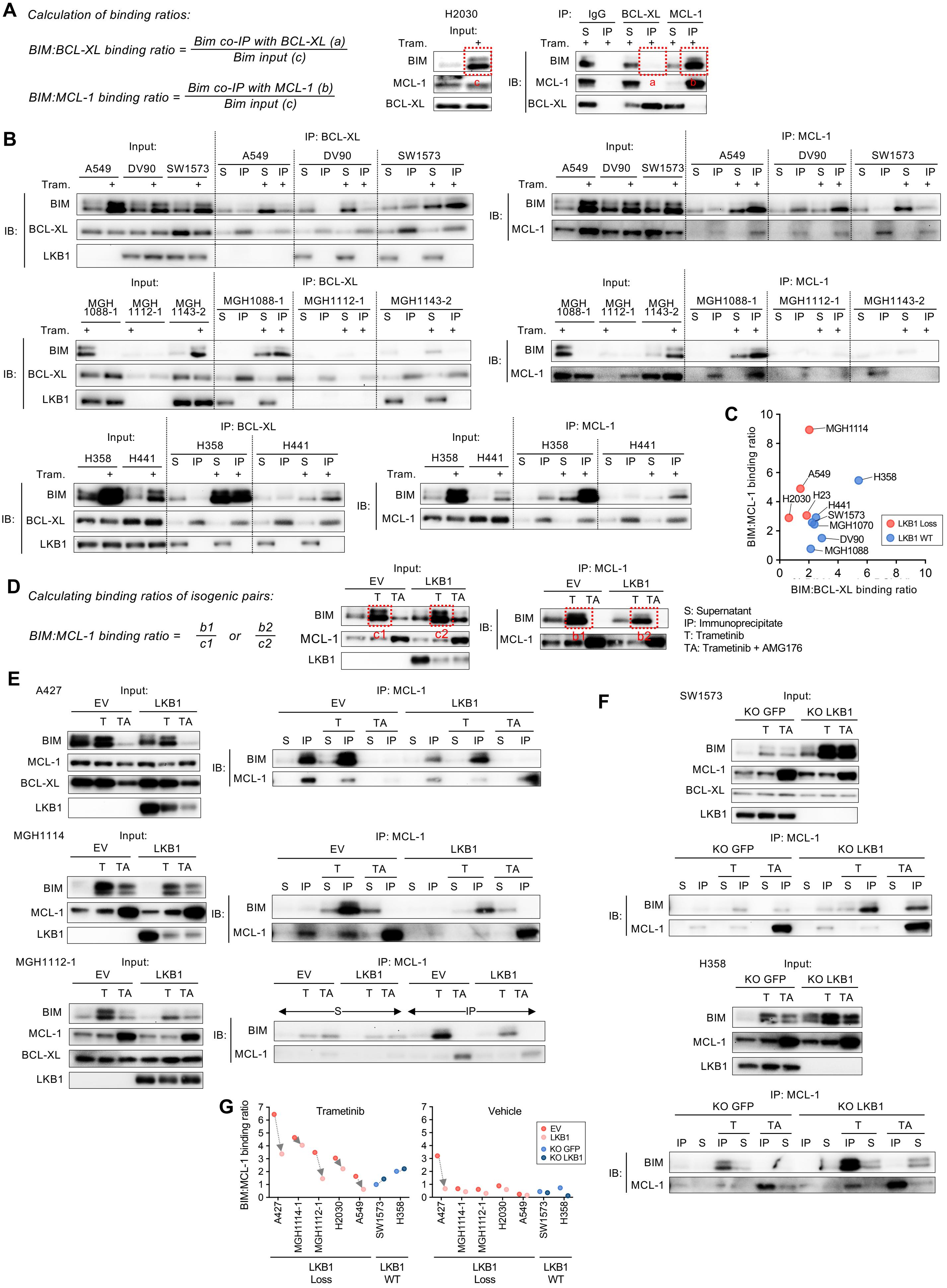


Supplemental Figure 8

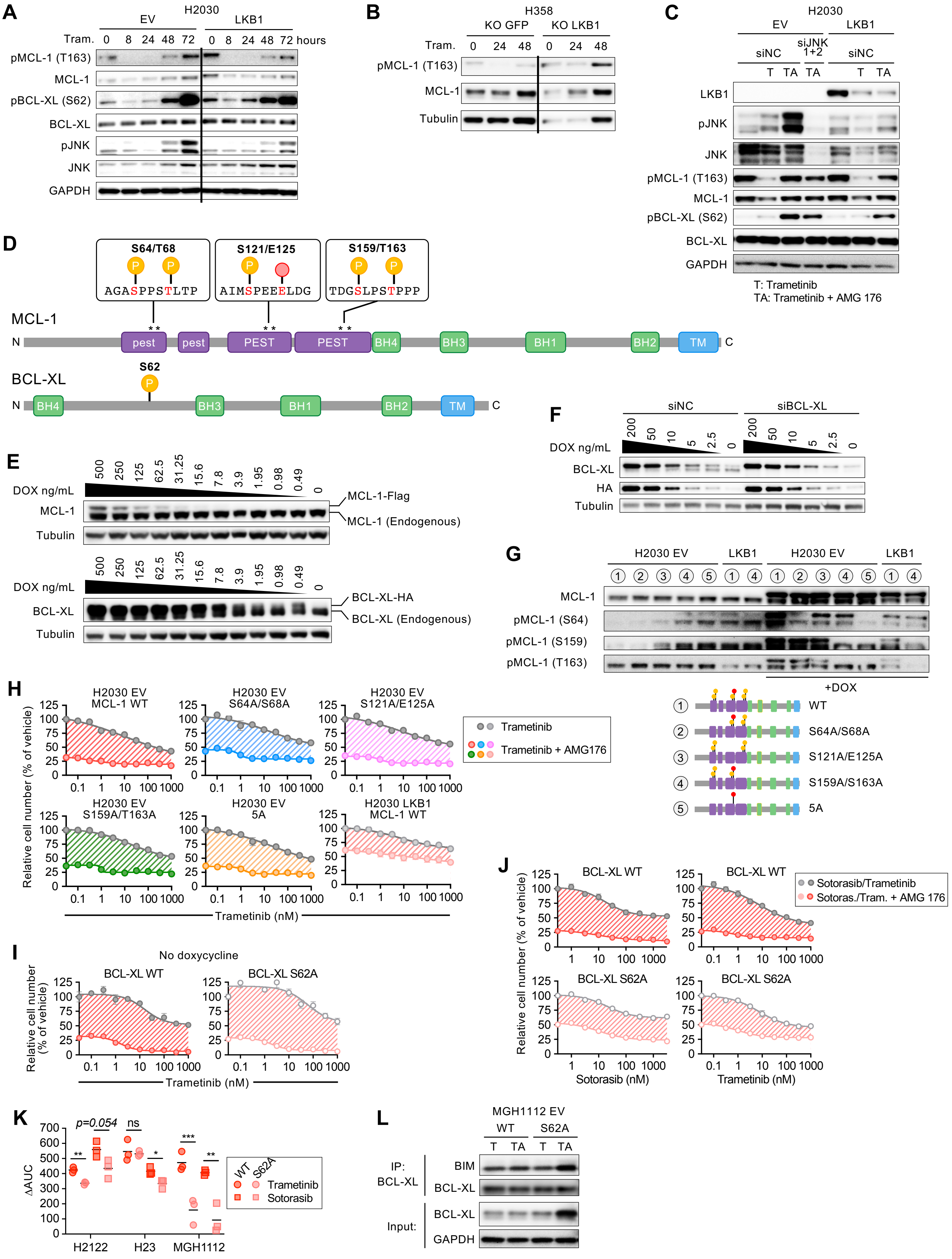


Supplemental Figure 9

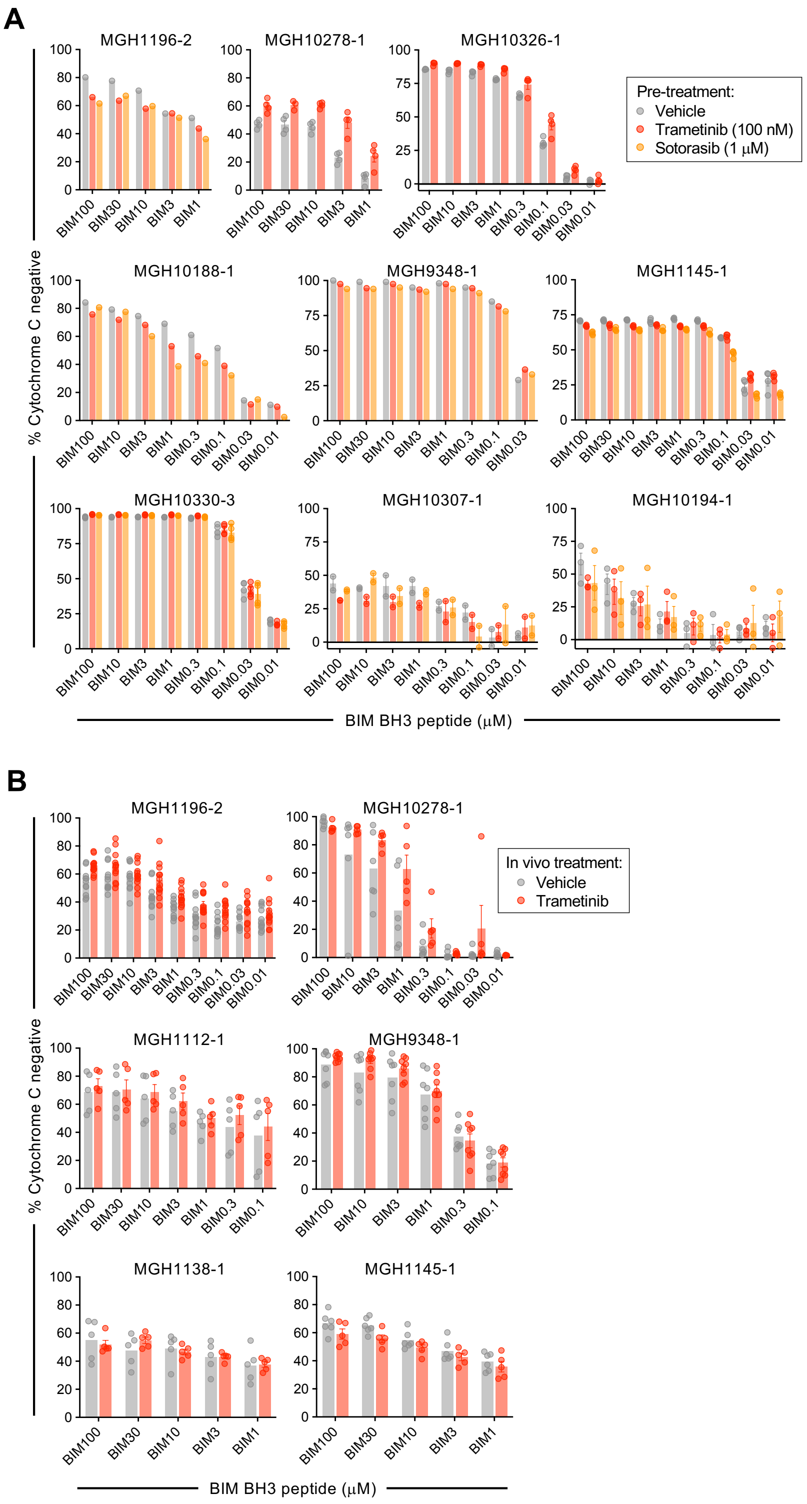


Supplemental Figure 10

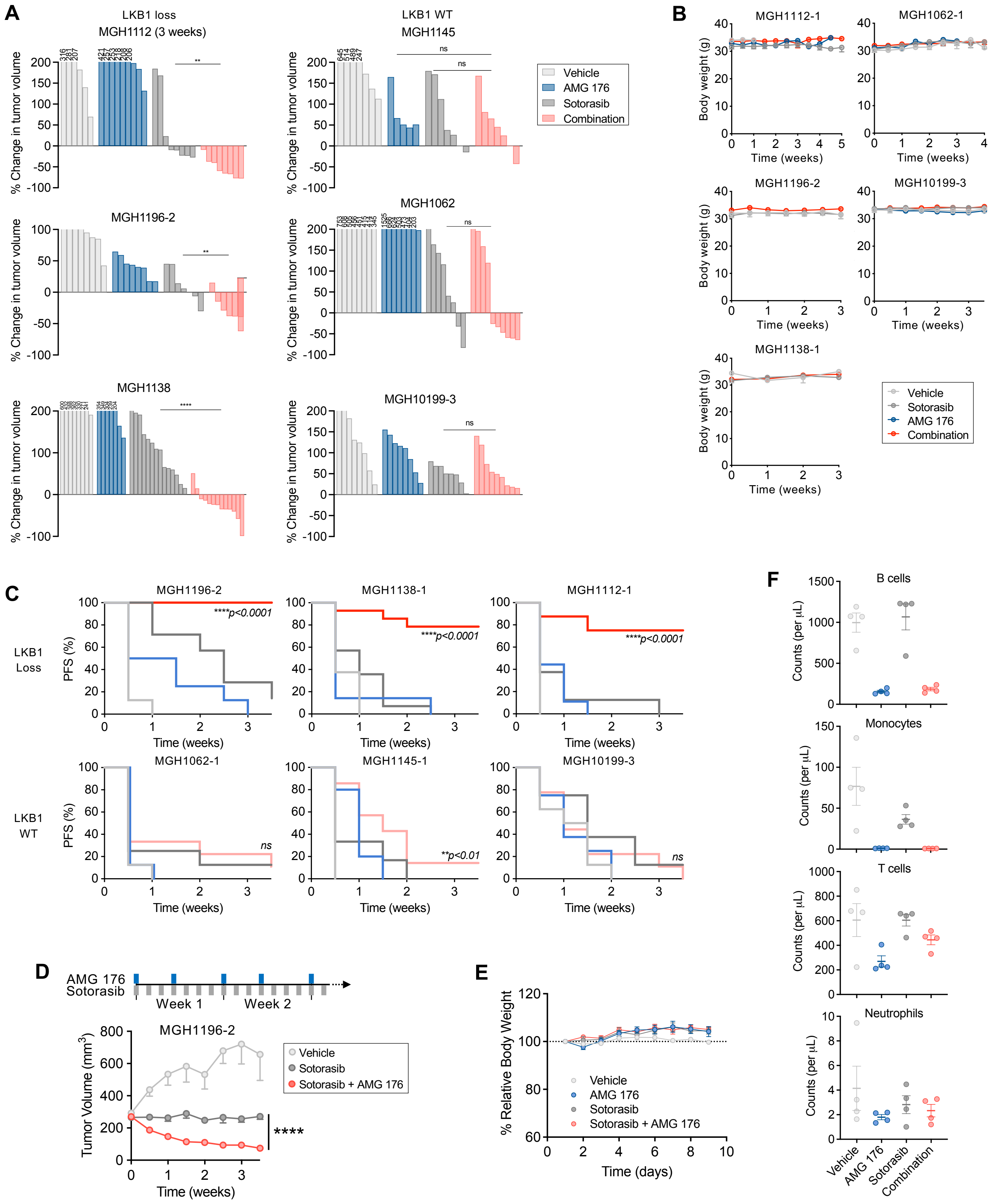


Supplemental Figure 11
